# Spinal neurons innervating multiple local and distant motor pools

**DOI:** 10.1101/2021.06.03.446906

**Authors:** Remi Ronzano, Camille Lancelin, Gardave S. Bhumbra, Robert M. Brownstone, Marco Beato

## Abstract

Motor neurons control muscle contractions, and their recruitment by premotor circuits is tuned to produce accurate motor behaviours. To understand how these circuits coordinate movement across and between joints, it is necessary to understand whether spinal neurons pre-synaptic to motor pools, project to more than one motor neuron population. Here, we used modified rabies virus tracing in mice to investigate premotor INs projecting to synergist flexor or extensor motor neurons, as well as those projecting to antagonist pairs of muscles controlling the ankle joint. We show that similar proportions of premotor neurons diverge to agonist and antagonist motor pools. Divergent premotor neurons were seen throughout the spinal cord, with decreasing numbers but increasing proportion with distance from the hindlimb enlargement. In the cervical cord, divergent long descending propriospinal neurons were found in contralateral lamina VIII, had large somata, were excitatory, projected to both lumbar and cervical motoneurons, and were at least in part of the V0 class. We conclude that distributed spinal premotor neurons coordinate activity across multiple motor pools and that there are spinal neurons mediating co-contraction of antagonist muscles.

## Introduction

The spinal cord is ultimately responsible for organising movement by controlling the activation pattern of motor neurons (MNs), which in turn produce appropriate patterns of muscle contractions to produce limb movement. Across any single limb joint, there are fundamentally 3 types of control – or 3 “syllables of movement” – possible. The 3 basic syllables are: (1) changing a joint angle, (2) stiffening a joint, and (3) relaxing a joint. The concatenation of these syllables across joints within and between limbs ultimately produces behaviour (Brownstone, 2020; Wiltschko et al., 2015).

To change a joint angle, MNs innervating synergist muscle fibres are activated whilst those that innervate antagonist muscle fibres are inhibited. This “reciprocal inhibition” (Eccles, 1969; Eccles et al., 1956), is mediated locally by spinal interneurons (INs) throughout the spinal cord; this syllable has been fairly well characterised, with responsible neurons identified and classified (Zhang et al., 2014).

The other 2 syllables are less well studied, but it is clear that behavioural joint stiffening requires co-activation of motor neurons innervating antagonist muscle groups, while joint relaxation would require co-inhibition of these motor neurons. Co-contraction has been studied, and has largely thought to result from brain activity (Humphrey and Reed, 1983), whereas circuits mediating co-inhibition remain elusive. Since the spinal cord controls movement not only across single joints but throughout the body, it is natural to consider whether it contains the circuits necessary to produce these different syllables.

To identify whether these syllables are produced by spinal circuits, several questions can be asked: Does the spinal cord contain circuits that lead to co-activation or co-inhibition of different pools of MNs – either synergists or antagonists? Does each motor pool have its own dedicated population of premotor INs, and are these INs interconnected in such a way that they can produce contraction of different muscle groups? Or are there populations of INs that project to multiple motor pools in order to effect contraction (or relaxation) of multiple muscles? Indeed, INs that have activity in keeping with innervation of multiple synergists, leading to motor “primitives” or synergies (Bizzi and Cheung, 2013; Giszter, 2015; Hart and Giszter, 2010; Takei et al., 2017; Tresch and Jarc, 2009) have been identified.

Normal behaviours require coordination of syllables across joints between forelimbs and hindlimbs. This coordination relies on populations of propriospinal neurons projecting in either direction between the lumbar and cervical enlargements (Eidelberg et al., 1980; Giovanelli Barilari and Kuypers, 1969; Miller and van der Meché, 1976; Ruder et al., 2016). Long descending propriospinal neurons (LDPNs) were first proposed in cats and dogs more than a century ago (Sherrington and Laslett, 1903), and their existence has been confirmed in several other species including humans (Alstermark et al., 1987a, 1987b; Ballion et al., 2001; Brockett et al., 2013; Flynn et al., 2017; Giovanelli Barilari and Kuypers, 1969; Jankowska et al., 1974; Mitchell et al., 2016; Nathan et al., 1996; Ni et al., 2014; Reed et al., 2009; Ruder et al., 2016; Skinner et al., 1979). While LDPNs that establish disynaptic connections to lumbar MNs were identified, it was initially suggested that at least some cervical LDPNs could establish monosynaptic inputs to lumbar MNs (Jankowska et al., 1974). This connectivity was later confirmed using monosynaptic modified rabies virus (RabV) tracing (Ni et al., 2014). More recently, descending and ascending spinal neurons and their involvement in the control of stability and interlimb coordination have been characterized, but these studies did not directly focus on monosynaptic premotor circuits (Pocratsky et al., 2017; Ruder et al., 2016). It is likely that LDPNs function to ensure coordination between fore and hindlimbs, and they could be an important source of premotor input to MNs, providing a substrate for coordination between distant joints.

In the present study, we examine circuits underlying co-activation and co-inhibition in the spinal cord by assessing premotor neurons through the use of RabV tracing techniques (Ronzano et al., 2021; Ugolini, 1995; Wickersham et al., 2007). We used glycoprotein (G)-deleted RabV (ΔG-Rab), and supplied G to MNs through crossing ChAT-Cre mice with RΦGT mice (Ronzano et al., 2021; Takatoh et al., 2013). We injected ΔG-RabV tagged with two different fluorescent proteins into hindlimb muscle pairs of ChAT-Cre mice to retrogradely trace premotor circuits throughout the spinal cord. At the lumbar level, this method revealed apparent low rates of INs projecting to both MN pools targeted. As the distance from targeted MN pool to premotor INs increased, the density of infected premotor INs decreased. But the apparent rate of divergence to multiple pools was higher in thoracic and cervical regions than in the lumbar spinal cord. Interestingly, the extent of divergence throughout the spinal cord was similar whether injections were performed in flexor or extensor pairs, or in synergist or antagonist pairs of muscles. In addition, population of premotor LDPNs was identified in the cervical spinal cord. These neurons had a high rate of divergence and large somata, projected contralaterally, were excitatory, located in lamina VIII, and projected to cervical MNs as well as lumbar MNs. Together, these data show that the spinal cord contains premotor INs that project to multiple motor pools (including antagonists), and could thus form substrates for the fundamental syllables of movement.

## Results

### Lumbar premotor INs reveal similar divergence patterns to synergist and antagonist motor pools

Given evidence that INs are involved in motor synergies (Hart and Giszter, 2010; Levine et al., 2014; Takei et al., 2017; Takei and Seki, 2010), we would expect that there would be INs in the lumbar spinal cord that project to synergist motor pools. We thus first investigated whether such premotor INs could be infected with two RabVs, one from each muscle. We injected ΔG-Rab expressing eGFP or mCherry into synergist ankle extensors (LG and MG) or synergist ankle flexors (TA and PL) in ChAT-Cre;RΦGT mice (Ronzano et al., 2021). These mice selectively express the rabies G in cholinergic neurons (including MNs), providing the necessary glycoprotein for retrograde trans-synaptic transfer from infected MNs to premotor INs (Figure 1A). After 9 days, we visualized the distribution of premotor INs that expressed one or both fluorescent proteins, specifying the premotor INs that make synaptic contact with two motor pools as “divergent” premotor INs (Figure 1B). We found divergent premotor INs distributed bilaterally in the ventral quadrants and ipsilaterally in the dorsal quadrant of the spinal cord (Figure 1C-D), consistently across experiments (Figure S1A, Table S3). Across the lumbar spinal cord, we quantified infected neurons on one of every three sections and found 4.0 ± 0.3 % (276/7043, n=4, 2 extensor and 2 flexor pairs) of labelled premotor INs were double-labelled, confirming that INs can be infected by more than one RabV. We would expect this to be an underestimate of INs that have divergent projections since RabV is not expected to label 100% of presynaptic neurons and as there is a reduced efficiency of double infections compared to single infections (Ohara et al., 2009) (see Discussion).

**Figure 1:**
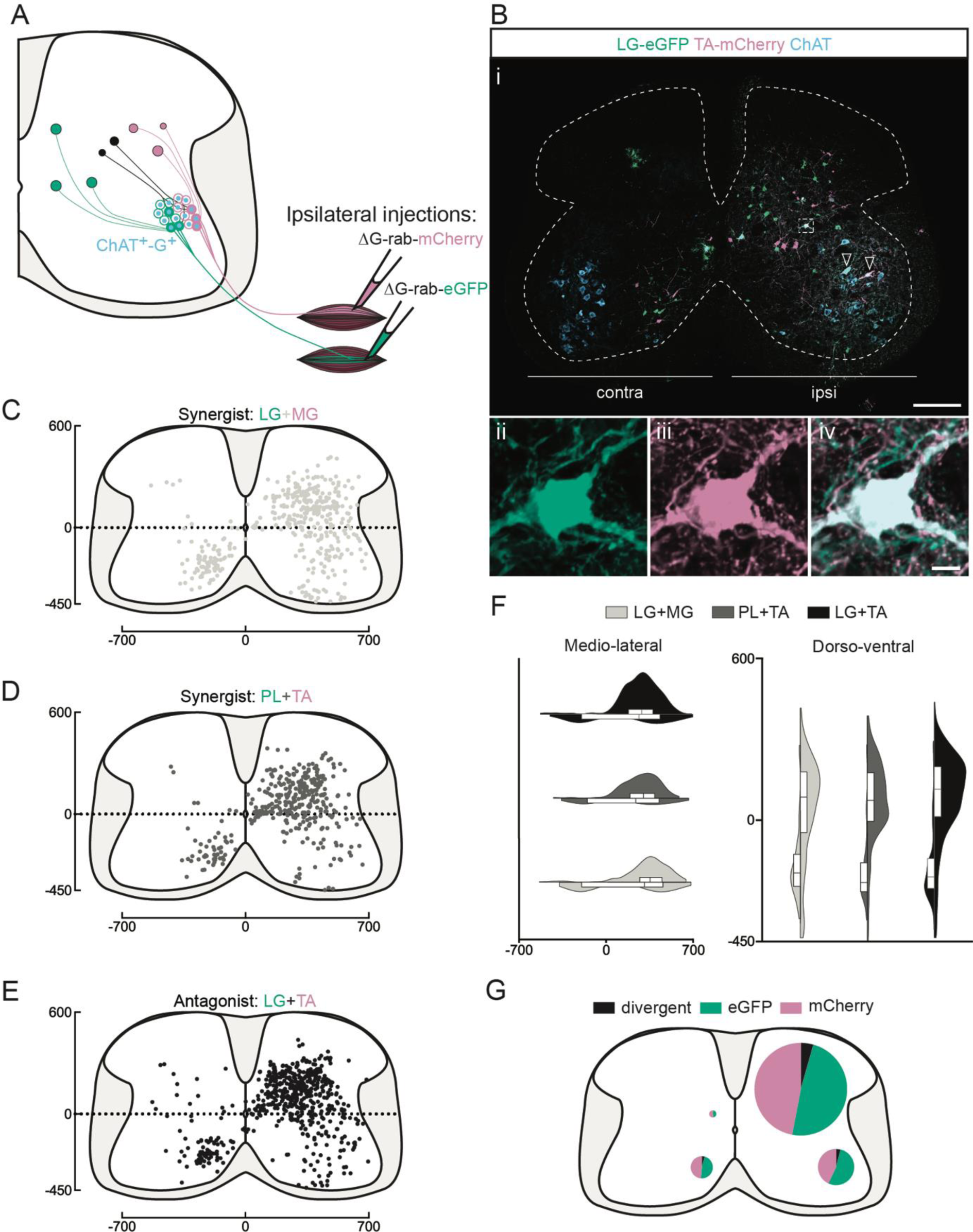
Divergent premotor INs organisation in the lumbar spinal cord. **(A)** Experimental strategy to describe divergent premotor INs that project to 2 motor pools of synergist (injection in TA and PL or LG and MG) or antagonist (TA and LG) pair of muscles. **(Bi)** Representative example of a lumbar transverse section following an injection in the TA (ΔG-Rab-mCherry) and LG (ΔG-Rab-eGFP), showing ChAT (grey blue), GFP (green) and mCherry (pink). A divergent premotor IN is highlighted in the dashed box. Arrowheads show infected MNs. The dashed contour drawn follows the grey matter contour. **(ii-iv)** Higher magnification of a divergent premotor IN that has been infected by the ΔG-Rab-eGFP and ΔG-Rab-mCherry. **(C-E)** Distribution of the lumbar divergent premotor INs following injections in (C) LG and MG (n=2), (D) PL and TA (n=2), (E) LG and TA (n=3). (**F**) Asymmetric violin plots showing the medio-lateral and dorso-ventral distributions of divergent premotor INs. The halves correspond respectively to the dorsal (top) and ventral (bottom) distributions and to the ipsilateral (right) and contralateral (left) distributions of divergent premotor INs in the lumbar cord. Violin areas were normalized on the number of divergent INs. **(G)** Distribution of the premotor INs within each quadrant of the lumbar cord, with pie sizes proportional to the percentage of premotor INs in each quadrant of the lumbar cord. Scale bars: (Bi) 200 µm; (Biv) 10 µm.

We next sought to determine whether this divergence was restricted to synergist motor pools or whether there are also premotor INs that diverge to antagonist pools and could thus be involved in co-contraction or joint stiffening. Following injections into flexor (TA) and extensor (LG) muscles, we found the rate of divergent premotor INs to be similar to that observed between synergist muscles, with 4.7 ± 0.5 % (206/4341, n=3 antagonist pairs, Figure 1E) double-labelled. The mapping of all divergent INs in every section revealed that, whether injections were in synergist (n=4, 2 extensor and 2 flexor synergist pairs) or antagonist (n=3 pairs) pairs of muscles, double-labelled premotor INs were distributed similarly (Figure 1F, 1G, Figure S1A, and Table S3 for summary of individual experiments). The proportion of divergent cells was calculated from the ratio of double and single infected cells in 1/3 sections, in order to avoid double counting cells present in consecutive sections (see Methods). Equal proportions of divergent premotor INs were found in the ventral ipsilateral quadrant (synergists: 74/1913 (3.9%) vs antagonists: (46/1046 (4.4%)), ventral contralateral quadrant (46/1020 (3.8%) vs 21/502 (4.2%)), and dorsal ipsilateral quadrant (153/3874 (3.9%) vs 134/2651 (5.1%)). There were few labelled neurons in the dorsal contralateral quadrant following either synergist or antagonist injections and a similarly low proportion were double labelled (in 1/3 sections: 3/236 (1.3%) and 5/142 (3.5%), respectively). Divergence in premotor circuits is thus common, with at least 1/25 (see Discussion) premotor INs diverging to 2 MN pools, whether synergists or antagonists.

Since motor synergies can span across more than a single joint, it is possible that divergent premotor INs could project to motor pools other than those injected. Indeed, following injection of ΔG-Rab-mCherry into the TA muscle, we could visualize mCherry positive excitatory (vGluT2+) boutons in apposition to L1 (Figure S2A-C), as well as to thoracic (as rostral as at least T10) MNs (Figure S2D,E), i.e., 3-7 segments rostral to the infected motor pool. This observation, in agreement with a previous study that described premotor INs coordinating the activity of multiple lumbar motor groups from L2 to L5 (Levine et al., 2014), supports the possibility that thoraco-lumbar premotor circuits comprise a substrate for multi-joint synergies.

### Thoracic premotor neurons project to multiple lumbar motor pools

In order to maintain posture and stability, trunk muscles are coordinated with hindlimb movements. Neurons in the thoracic cord that are premotor to lumbar MNs have previously been described (Ni et al., 2014); we thus next examined the projections of thoracic premotor neurons to lumbar motor pools. These premotor neurons were found with decreasing density from T11 through T2 whether the injections were in flexor (TA and PL; Figure 2A) or extensor (LG and MG) pairs of muscles. The distributions of single labelled as well as divergent premotor neurons were similar whether injections were performed in flexor, extensor, or antagonist pairs of muscles (Figure 2 B-F). Divergence rates calculated from the whole thoracic spinal cords were similar between synergist and antagonist injections with 16.2 ± 5.7% (77/497, n=4, 2 extensor and 2 flexor pairs, Figure 2B, C) and 9.0 ± 0.7% (59/401, n=3 antagonist pairs, Figure 2D), respectively. In all animals (7/7), the overall proportion of double-labelled neurons in the thoracic spinal cord was higher than in the lumbar cord (13.1 ± 5.6%, Figure 7B).

**Figure 2:**
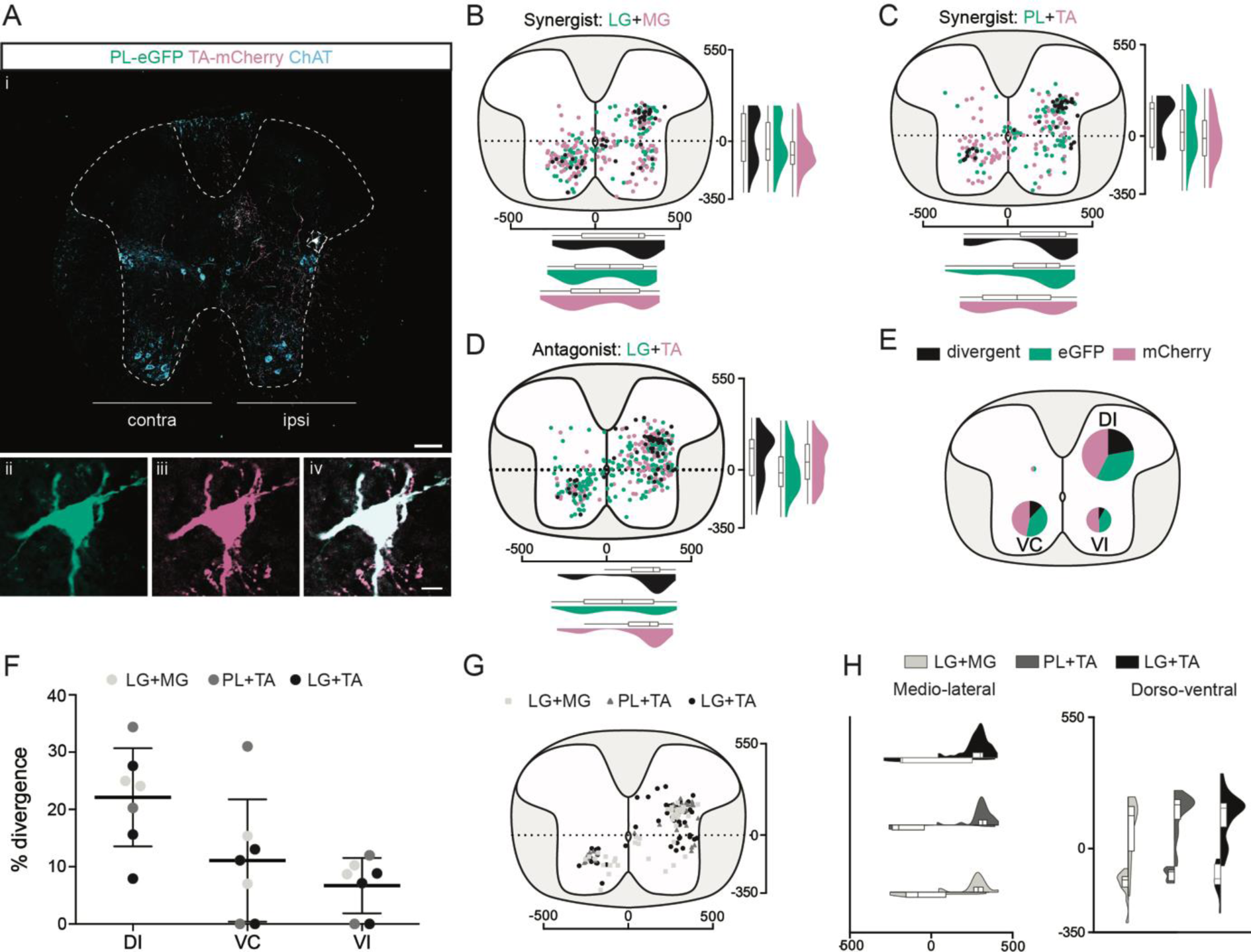
Divergent premotor neurons organisation in the thoracic part. **(Ai)** Representative example of a thoracic transverse section following an injection in the PL (ΔG-Rab-eGFP) and TA (ΔG-Rab-mCherry), showing ChAT (grey blue), GFP (green) and mCherry (pink). A divergent premotor neuron is highlighted in the dashed box. The dashed contour drawn follows the grey matter contour. **(ii-iv)** Higher magnification of a divergent premotor neuron that has been infected by both ΔG-Rab-eGFP and ΔG-Rab-mCherry. **(B-D)** Distribution of the thoracic premotor neurons infected following injections in (B) LG and MG (n=2), (C) PL and TA (n=2) and (D) LG and TA (n=3). Divergent premotor neurons infected from both injections are labelled in black. The violin plots show the dorso-ventral and medio-lateral distributions of divergent (black), GFP positive (green) and mCherry positive (pink) premotor neurons along the medio-lateral and dorso-ventral axis. Each violin area is normalised to 1. **(E)** Pies showing the distribution of infected premotor neurons in each quadrant; the size of the pies is proportional to the number of infected neurons. **(F)** Plot showing the divergence rate in each quadrant of the thoracic cord. DI: dorsal ipsilateral; VC: ventral contralateral; VI: ventral ipsilateral. **(G)** Overlap of distributions of divergent thoracic premotor neurons followings each pair of muscles injected. (**H**) Asymmetric violin plots showing the medio-lateral and dorso-ventral distributions of divergent premotor neurons. The halves correspond respectively to the dorsal (top) and ventral (bottom) distributions and to the ipsilateral (right) and contralateral (left) distributions of divergent premotor neurons in the thoracic cord. Violin areas were normalized on the number of divergent neurons. Scale bars: (Ai) 100 µm, (Aiv) 10 µm.

In all animals (7/7), the majority of divergent premotor neurons in the thoracic cord were located in the ipsilateral dorsal quadrant (46/77, n=4 synergist and 42/59, n=3 antagonist pairs, Figure 2 E-H), with 22.1 ± 8.6% double labelled (46/188 synergists and 42/211 antagonists, Figure 2G-H). The divergence rates in the two ventral quadrants were lower: in the ventral cord, double-labelled neurons were observed in 5/7 animals (3/4 synergist; 2/3 antagonist in both quadrants) ipsilaterally (6.7 ± 4.8%; 10/118 synergist and 5/70 antagonist pairs), as well as contralaterally (11.1 ± 10.7%; 21/167 synergist and 12/104 antagonist pairs) to the injection (Figure 2G). Thus, there are premotor neurons throughout the thoracic cord that project directly to more than one motor pool, including antagonist pairs, in the lumbar spinal cord, with most of these located in the ipsilateral dorsal quadrant.

### Cervical premotor long propriospinal descending neurons diverge and share a typical location and morphology

Cervical long descending propriospinal neurons (LDPNs) have been shown to modulate interlimb coordination to provide stability (Eidelberg et al., 1980; Miller and van der Meché, 1976; Ruder et al., 2016). Given that cervical premotor LDPNs projecting to TA MNs have previously been demonstrated (Ni et al., 2014), we asked whether these neurons could be premotor to hindlimb and/or hindlimb-forelimbs MN pairs.

We found that premotor LDPNs projecting to flexor (TA and PL) and extensor (LG and MG) MNs were localised throughout the rostrocaudal extent of the ventral cervical cord. Of 92 premotor LDPNs, 88 were localized in the ventral quadrants, 68 of which were in contralateral lamina VIII (n=7, 4 synergist and 3 antagonist pairs, Figure 3A-G). A substantial proportion of premotor LDPNs was double-labelled, with the proportion and location of double-labelling similar across experiments (Figure S1B and Table S3) whether injections were into synergist or antagonist pairs (42.4 ± 22.1 % per animal, total of 19/55 neurons, n=4 synergist pairs and 47.9 ± 7.1 % per animal, total of 19/37 neurons, n=3 antagonist pairs, Figure 3E-F). This apparent divergence rate of LDPNs in the cervical cord was higher than in the lumbar and thoracic cords in all animals (7/7, Figure 7B). These divergent premotor LDPNs exhibited a stereotypical morphology with an unusually large soma (774 ± 231 µm^2^, n=38 premotor LDPNs) compared to the double-labelled premotor neurons in the thoracic and lumbar cords (respectively 359 ± 144 µm^2^ and 320 ± 114 µm^2^, n=135 premotor neurons (thoracic), n=61 premotor INs (lumbar), p<0.0001 Kruskal-Wallis test, p<0.0001 (lumbar vs cervical) and p<0.0001 (thoracic vs cervical), Dunn’s multiple comparisons test). On average, the cross-sectional area of divergent cervical LDPNs was comparable to that of cervical MNs (661 ± 86 µm^2^, n=17 MNs, Figure 3H). Their location and size suggests that these divergent, commissural cervical premotor LDPNs may constitute a somewhat homogeneous population.

**Figure 3:**
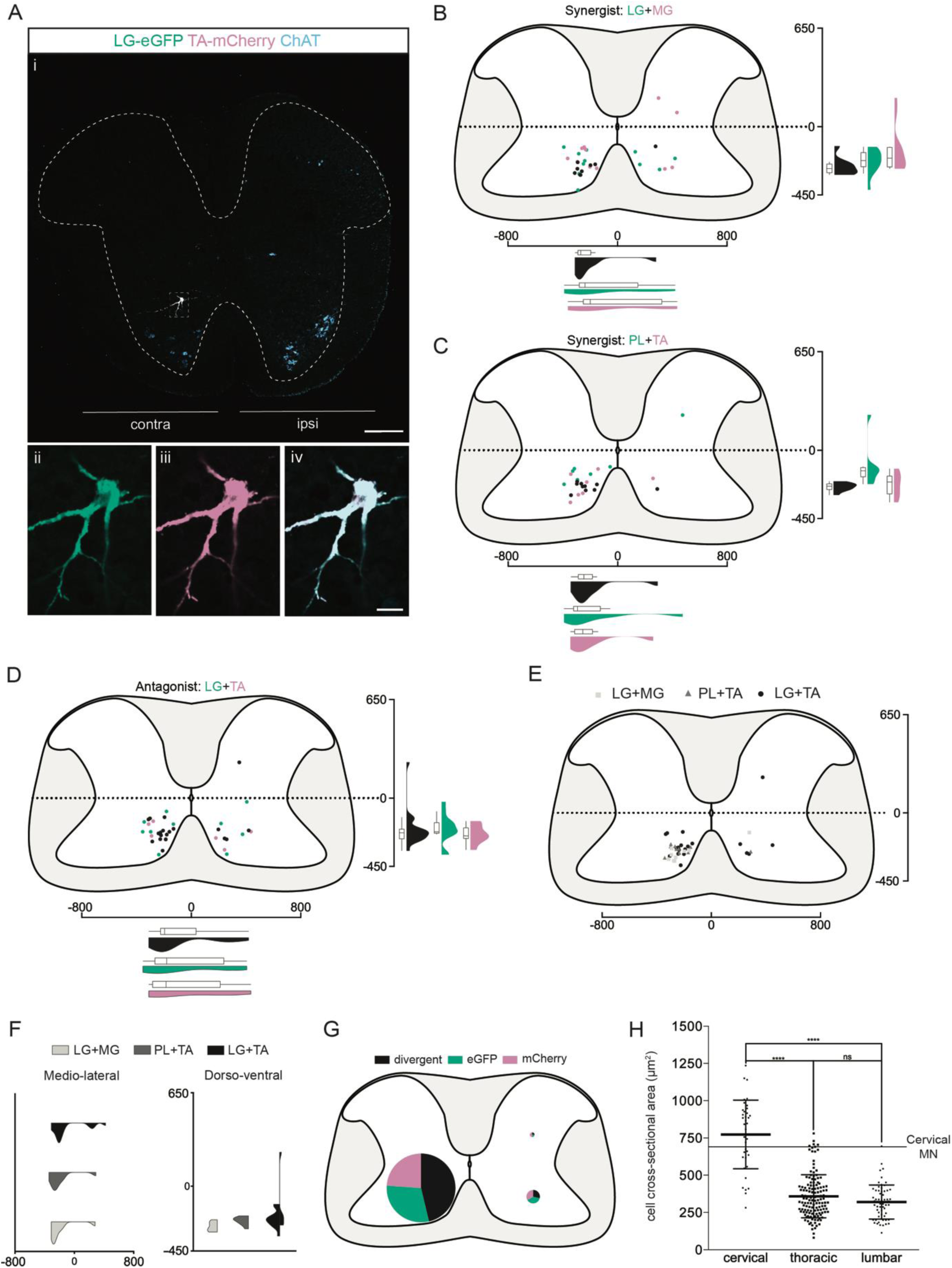
Divergent premotor LDPNs organisation in the cervical part. **(Ai)** Representative example of an upper cervical transverse section following an injection in the LG (ΔG-Rab-eGFP) and TA (ΔG-Rab-mCherry), showing ChAT (grey blue), GFP (green) and mCherry (pink). A divergent premotor LDPN is highlighted in the dashed box. The dashed contour drawn follows the grey matter contour. **(ii-iv)** Higher magnification of the divergent premotor LDPN. **(B-D)** Distribution of the cervical premotor LDPNs following injections in (C) LG and MG (n=2), (D) PL and TA (n=2) and (E) LG and TA (n=3). Divergent premotor LDPNs infected from both injections are labelled in black. The violin plots show the dorso-ventral and medio-lateral distributions of divergent (black), GFP positive (green) and mCherry positive (pink) premotor LDPNs along the medio-lateral and dorso-ventral axis. Each violin area is normalised to 1. **(E)** Overlap of the distribution of cervical divergent premotor LDPNs followings each pair of muscles injected. **(F)** Asymmetric violin plots showing the medio-lateral and dorso-ventral distributions of premotor divergent LDPNs. The halves correspond respectively to the dorsal (top) and ventral (bottom) distributions and to the ipsilateral (right) and contralateral (left) distributions of divergent premotor LDPNs in the cervical cord. Violin areas were normalized on the number of divergent neurons. (**G**) Pies showing the distribution of infected premotor LDPNs in each quadrant; the size of the pies is proportional to the number of infected premotor LDPNs in each quadrant. **(H)** Plot showing the distribution of the sectional areas of divergent premotor neurons in each part of the cord. The dashed line (labelled cervical MN) corresponds to the mean sectional area of cervical MNs (n=17 MNs). Scale bars: (Ai) 200 µm; (Aiv) 20 µm.

### Cervical premotor LDPNs are primarily excitatory

To determine the neurotransmitter phenotype of the premotor LDPNs, we used single ΔG-Rab-mCherry injections in ChAT-Cre;RΦGT mice crossed with mice expressing eGFP under the control of the promoter for the neuronal glycine transporter GlyT2 (Zeilhofer et al., 2005), Figure 4A). GlyT2 is expressed in the vast majority of spinal inhibitory INs (Todd et al., 1996; Todd and Sullivan, 1990) making GlyT2-eGFP mice a suitable tool to determine whether premotor LDPNs are inhibitory. Given that at least 40% of the labelled INs in the cervical region are divergent (see above), many of the neurons labelled following even single RabV injections would be expected to be divergent. Following injection into LG (Figure 4A), we found that only 1/21 infected cervical commissural premotor LDPNs was eGFP positive (n=3 LG injections, Figure 4B,C,F). Since none of the labelled neurons expressed ChAT, the majority of cervical premotor LDPNs are likely to be glutamatergic by exclusion. However, in agreement with previous results from TA injections (Ni et al., 2014), single-labelled thoracic premotor neurons comprised a mixed population of inhibitory and non-inhibitory neurons (34.4 ± 5.9%, 96/273, mCherry+ eGFP+ premotor neurons, n=3 LG injections, Figure 4D-F). Therefore, the neurotransmitter phenotype of divergent thoracic premotor neurons could not be determined.

**Figure 4:**
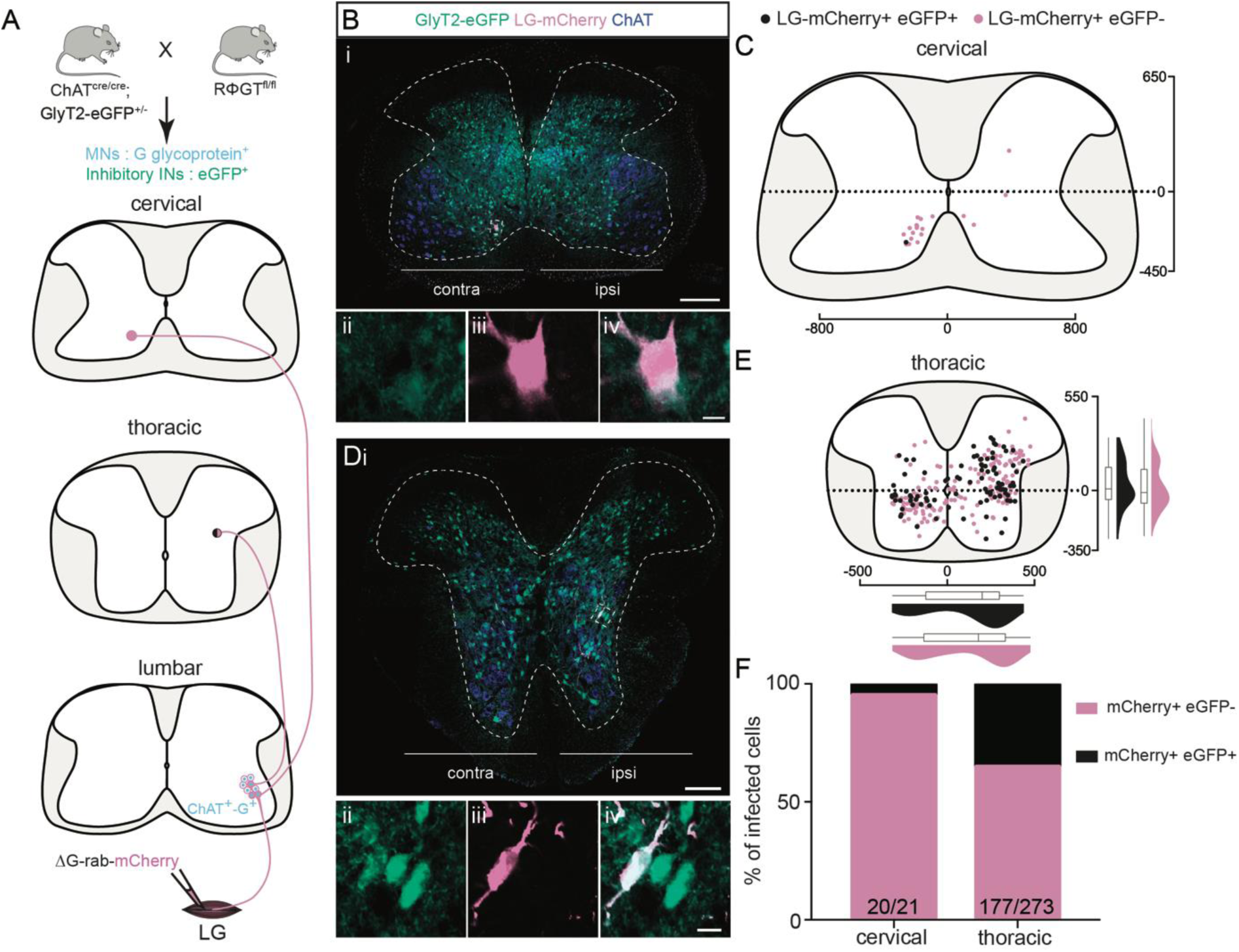
Non-inhibitory cervical premotor LDPNs and mixed populations of inhibitory and non-inhibitory thoracic premotor neurons revealed by injections in GlyT2-eGFP; RΦGT mice. **(A)** Experimental strategy to determine whether thoracic and cervical premotor neurons are inhibitory. **(B, D)** Representative example of (Bi) a cervical and (Di) a thoracic transverse section following an injection in the LG (ΔG-Rab-mCherry) using GlyT2-eGFP; RΦGT mice, showing ChAT (blue), GFP (green) and mCherry (pink). The dashed boxes highlight the infected premotor LDPNs. The dashed contours drawn follows the grey matter contours. **(Bii-iv, Dii-iv)** Higher magnification of the dashed box areas, highlighting (Bii-iv) a GFP-, mCherry+ cervical premotor LDPN on the contralateral Lamina VIII and (Dii-iv) a GFP+, mCherry+ thoracic premotor neuron in ipsilateral intermediate lamina. **(C, E)** Distribution of the (C) cervical and (E) thoracic premotor neurons infected, following injections in the LG of GlyT2-eGFP; RΦGT mice (n=3). The violin plots show the dorso-ventral and medio-lateral distributions of GFP+, mCherry+ (black) and GFP+, mCherry- (pink) premotor neurons along the medio-lateral and dorso-ventral axis. Each violin area is normalised to 1. **(F)** Proportions of inhibitory premotor neurons in the thoracic and the cervical part of GlyT2-eGFP; RΦGT mice following injections in the LG (n=3). Scale bars: (Bi) 200 µm; (Di) 100 µm; (Biv,Div) 10 µm.

### Cervical premotor LDPNs likely arise from the V0v domain

We next sought to determine the genetic provenance of divergent cervical LDPNs. Among the classes of interneurons defined by the early expression of transcription factors (Lee and Pfaff, 2001), the V0 and V3 cardinal classes are known to project to contralateral MNs. Of these, subclasses of V0 and all V3 INs are excitatory. These classes can be further subdivided, with all V3 subclasses being glutamatergic (Zhang et al., 2008), and V0 INs being neuromodulatory (V0_C_, cholinergic, (Miles et al., 2007), inhibitory (V0_D_, dorsal, Talpalar et al., 2013), or excitatory (V0_V_, ventral, (Talpalar et al., 2013), or V0_G_, medial glutamatergic neurons that project to dorsal and intermediate lamina but not to MNs, (Zagoraiou et al., 2009)). Since previous studies showed that none of the LDPNs with soma in the cervical cord belong to the V3 population (Flynn et al., 2017), we focussed on determining whether these LDPNs were of the V0 class.

V0 INs are defined by their embryonic expression of the transcription factor Dbx1 (Alaynick et al., 2011). However, Dbx1 cannot reliably be detected in postnatal cords at the age of our mice. On the other hand, Lhx1 is expressed throughout the V0 and V1 populations and may be detectable at this early postnatal stage (Skarlatou et al, 2020). Since V1 and V0_D_ INs are glycinergic (Alvarez et al, 2013; Talpalar et al, 2013; Lanuza et al, 2004), and V0_C_ are cholinergic (Miles et al, 2007), and since we have shown that LDPNs are negative for GlyT2 and ChAT, we interpreted the expression of Lhx1 as evidence of the cervical LDPNs belonging to the V0_V_ class. Following injection of gastrocnemius (GS, n=4, Figure 5A), we detected 33 premotor LDPNs. Of these infected cervical premotor LDPNs, 8 (∼24%) were clearly Lhx1 positive (Figure 5B, 5C). Given that there is a decrease of Lhx1 expression along the course of postnatal development (Figure S3), the proportion of premotor LDPNs that were positive for Lhx1 expression was likely underestimated. Thus, at least some of the divergent premotor LDPNs arise from V0_v_ neurons.

**Figure 5:**
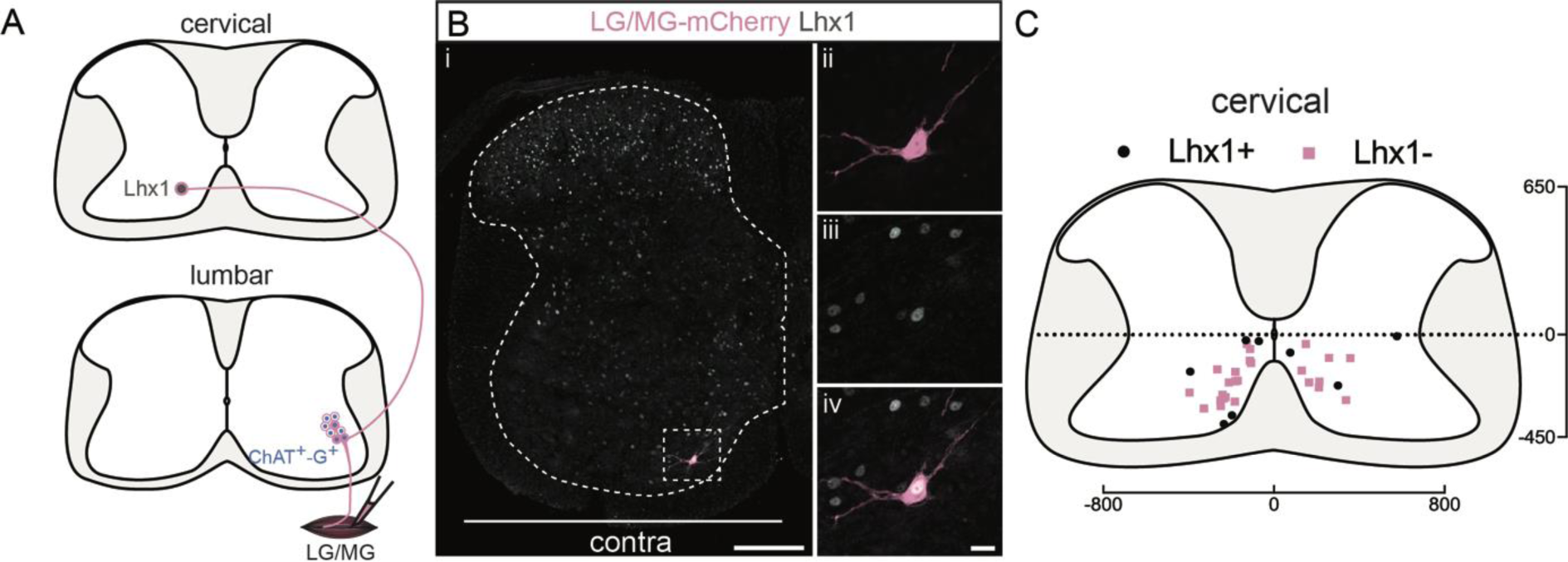
A subpopulation of cervical premotor LDPNs expresses Lhx1. **(A)** Experimental strategy to determine whether cervical premotor LDPNs express Lhx1. (**Bi**) Representative example of a transverse section from the cervical cord following an injection in GS (ΔG-Rab-eGFP) muscles, showing a cervical premotor LDPN infected (pink) expressing Lhx1. The premotor LDPN expressing Lhx1 is highlighted in the dashed box. The dashed contour drawn follows the grey matter contour. (**ii-iv**) Higher magnification of the premotor LDPN Lhx1+ that has been infected by the ΔG-Rab-mCherry. (**C**) Distribution of the cervical premotor LDPNs following injections in GS whether they are Lhx1+ (black) or not (pink). Scale bars: (Bi) 200 µm; (Biv) 20 µm.

### Cervical premotor LDPNs also project to local cervical MNs

Given that propriospinal neurons are involved in interlimb coordination, we next sought to determine whether the divergent cervical premotor LDPNs also project to cervical MNs. We therefore performed a series of experiments in which we injected forearm muscles (FMs; Table S2) with ΔG-Rab-mCherry, and extensor hindlimb GS with ΔG-Rab-eGFP. Given the involvement of LPDNs in the diagonal synchronisation of fore and hindlimb during locomotion (Bellardita and Kiehn, 2015; Ruder et al., 2016; Sherrington et al., 1906), we first injected contralateral FMs and GS (Figure S4A). We found that 2/26 cervical premotor LDPNs were also infected from the FMs injection with 1 divergent LDPNs in the lamina VIII contralateral to the hindlimb injection in each of 2 of the 3 injected animals (Figure S4B). Thus, at least a few cervical premotor LDPNs monosynaptically project to diagonal lumbar and cervical MNs. However, given the paucity of these cervical premotor LDPNs projecting to local cervical MNs, we could not reliably determine whether this subpopulation shared the same morphology as described above.

As it has also been suggested that LDPNs participate in ipsilateral control of fore- and hindlimb (Miller and van der Meché, 1976), we sought to determine if premotor LDPNs project to homolateral lumbar and cervical motor pools (Figure 6A). When homolateral limbs were targeted, we found that some premotor LDPNs infected from ankle extensor injections were also infected from homolateral FM injection (in 5/6 animals, 16/80 premotor LDPNs were also infected from FMs injection 18.7 ± 12.9%, Figure 6B-D). These divergent premotor LDPNs that projected to lumbar and cervical MNs were all located in the ventral quadrants with 11/16 located in contralateral lamina VIII, and were distributed throughout the rostrocaudal extent of the cervical cord, including segments rostral (C4) to the MN pools innervating the injected forelimb muscles. Furthermore, they had a soma size similar to the premotor LDPNs double labelled by dual hindlimb injections (632 ± 236 µm^2^, p=0.056, n_1_=16 premotor LDPNs infected from both homolateral fore and hindlimb injections vs n_2_=38 divergent premotor LDPNs infected from dual hindlimb injections (see above), Mann-Whitney test; Figure 6E). This result indicates that these neurons constitute a somewhat homogenous subpopulation that at least in part diverges to more than one lumbar motor pool and at least one homolateral cervical motor pool.

**Figure 6:**
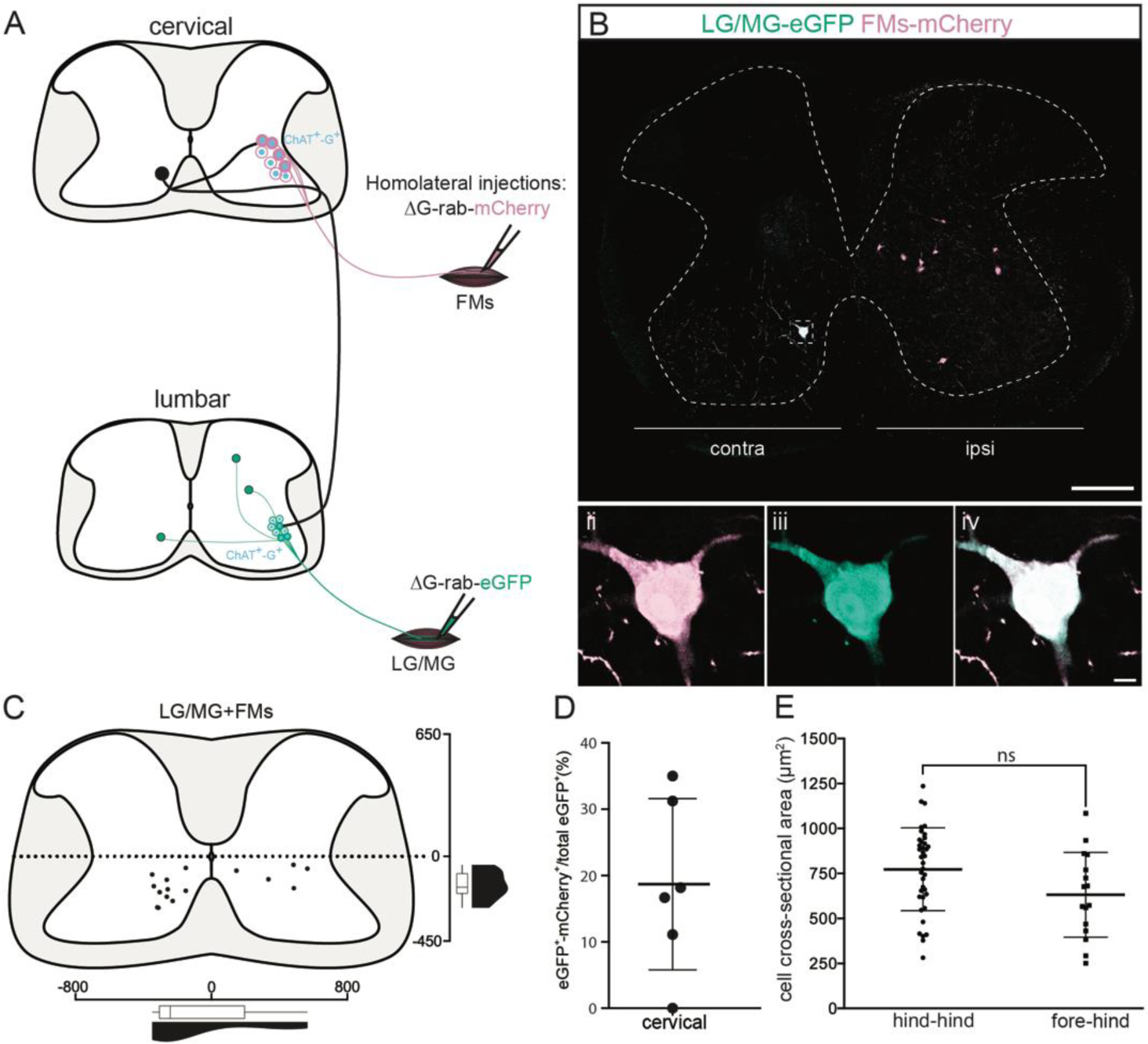
Cervical premotor LDPNs also innervate homolateral lumbar and cervical MNs. **(A)** Experimental strategy to determine whether divergent cervical premotor LDPNs that innervate homolateral lumbar and cervical MNs do exist. **(Bi**) Representative example of a transverse section from the cervical cord following an injection in FMs (ΔG-Rab-mCherry) and GS (ΔG-Rab-eGFP) muscles, showing a premotor LDPN infected from the two contralateral motor pools. The dashed box highlights the divergent premotor LDPN. The dashed contour drawn follows the grey matter contour. **(ii-iv)** Dashed box area at higher magnification. **(C)** Distribution of the premotor LDPNs infected from the homolateral injections in GS and FMs. The violin and box plots show the distribution of divergent premotor LDPNs innervating homolateral local FMs and distant GS motor pools along the medio-lateral and dorso-ventral axis. Each violin area is normalised to 1. **(D)** Proportion of cervical premotor LDPNs that also project to FM motor pools per animal. **(E)** Plot showing the sectional area of the cervical divergent premotor LDPNs that diverge to two pools of lumbar MNs (hind_hind) and to the pools of GS and FM MNs (fore_hind). Scale bars: (Bi) 200 µm; (Biv) 10 µm.

**Figure 7:**
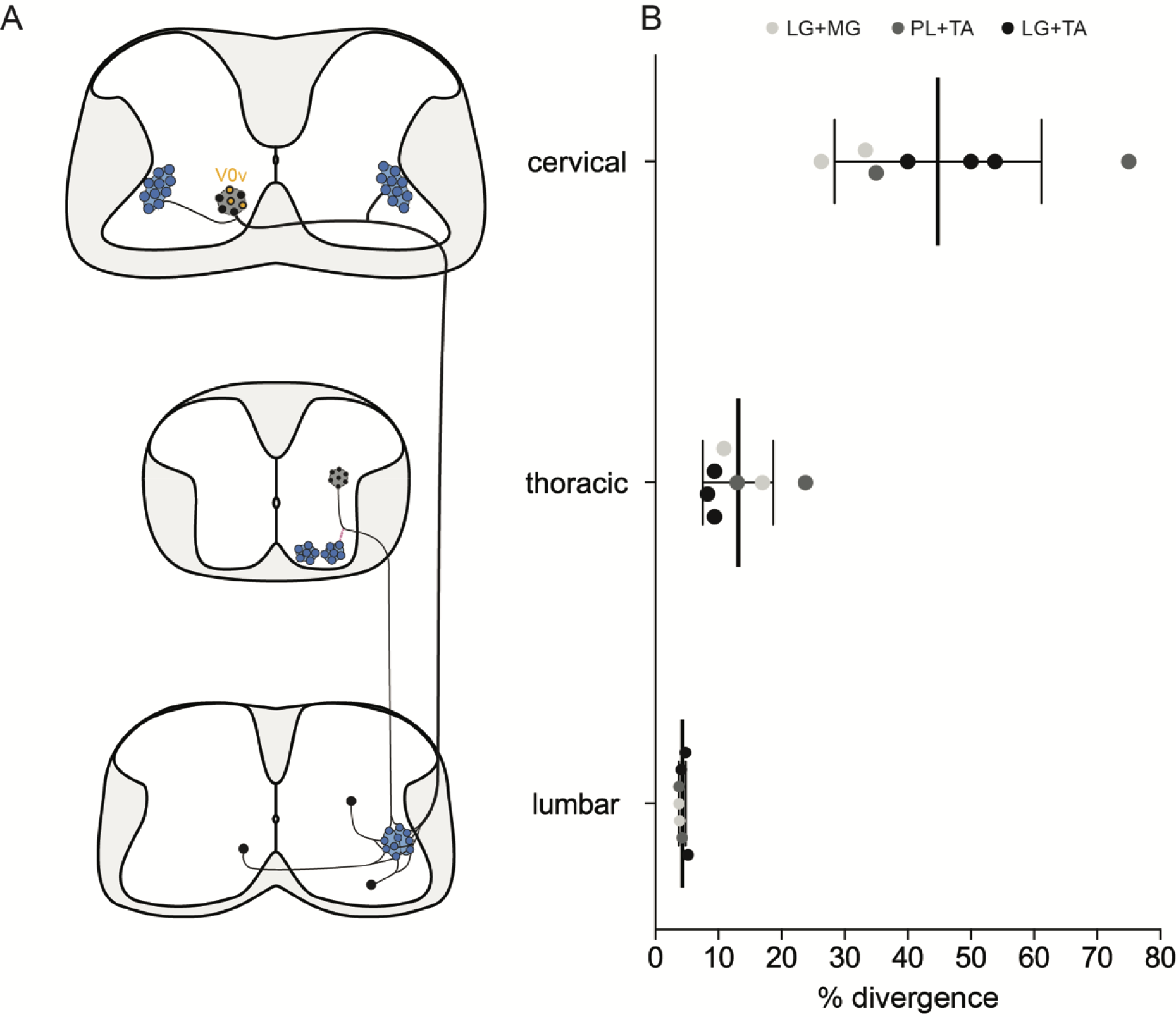
Divergence rates throughout the spinal cord and circuits unravelled. **(A)** Schematic summarizing the projections unravelled by the study. **(B)** Plot showing the increase of the apparent divergence rate with the distance between innervated MNs and premotor neurons.

### Distribution of premotor long ascending propriospinal neurons differs from that of LDPNs

Having identified a homogeneous population of divergent premotor long descending propriospinal neurons with projections from the cervical to the lumbar region, we next investigated whether ascending propriospinal neurons projecting from the lumbar or thoracic segments to cervical MNs could be identified. Following FMs injections, ascending premotor INs were observed throughout the cord (thoracic to sacral). There were very few (<1%) bifurcating (ascending/descending) premotor neurons in the thoracic cord after injections in homolateral GS and FMs (4/523 double labelled premotor neurons between T2 and T11, n=3, Figure S5A-B).

We identified premotor long ascending propriospinal neurons (LAPNs) in the lumbar cord, about half of which were localised in the dorsal ipsilateral quadrant (56/117, n=6 fore-hindlimb injections). This distribution indicates that lumbar premotor LAPNs are positioned differently from cervical premotor LDPNs, which were almost exclusively ventral (164/172, n=13 pair of injections, see above). Of the 117 lumbar premotor LAPNs identified, 10 were also labelled from GS injections, indicating that some neurons projected both to local lumbar MNs as well as to cervical MNs (n=6 ipsilateral fore-hindlimb injections, Figure S5C-D). However, the position of these divergent premotor LAPNs was different from that of the premotor LDPNs, in that they were distributed throughout the 4 quadrants of the cord (Figure S5D).

Finally, we turned our attention to the sacral spinal cord, where we found few premotor LAPNs (12 neurons in 4 of 6 mice). Of these, however, 10/12 were in the ventral contralateral quadrant (n=6 ipsilateral fore-hindlimb injections, Figure S5E-F), similar to the location of the cervical premotor LDPNs. Like these cervical neurons, the sacral LAPNs had strikingly large somata (710 ± 310 µm^2^, n=11 premotor LAPNs, Figure S5G). Of 12 labelled neurons, 3 were also infected from the hindlimb (LG) injections (Figure S5E-G). The size and location of these sacral premotor LAPNs were similar to the population of cervical divergent premotor LDPNs, and they may thus represent a “reverse counterpart” of this descending system.

## Discussion

Animals perform rich repertoires of movements through controlling muscle contractions around joints to produce the fundamental syllables of movement (Brownstone, 2020). To understand how behavioural repertoires are formed, it is important to understand the organization of the neural circuits underlying the production of each syllable. By using monosynaptic restricted RabV tracing techniques, we investigated the presence of spinal premotor circuits underlying co-activation (joint stiffening) or co-inhibition (joint relaxation) of motor pools across joints and between limbs. We found that at least 1/25 local lumbar premotor INs projects to multiple motor pools, in similar proportions whether these pools were synergist or antagonist pairs. Furthermore, we found that whereas the density of premotor neurons decreases with distance rostral to the motor pool targeted, a high proportion of labelled cervical LDPNs projects to multiple motor pools. These premotor LDPNs are in contralateral lamina VIII, had large somata, are excitatory, and project to multiple motor pools including those in the lumbar and cervical enlargements. These divergent neurons could thus form a substrate for joint and multi-joint stiffening that contributes to the production of a fundamental syllable of movement.

### Estimating proportions of divergent premotor interneurons

The control of MNs across motor pools through spinal premotor circuits is required for the performance of all motor tasks involving limb movements. Previous studies showed the importance of motor synergies in the production of complex movements (Giszter, 2015; Takei et al., 2017), with the spinal cord identified as a potential site for muscle synergy organisation (Bizzi and Cheung, 2013; Levine et al., 2014). In this regard, it might be expected that a significant proportion of local spinal premotor INs innervate multiple motor pools, in particular those corresponding to synergist muscles. Perhaps surprisingly, we found similar rate of divergence throughout the spinal cord be the targeted MN pools synergist or antagonist. In the lumbar part, at least 4% of the local premotor INs project to two motor pools. Distally in the thoracic as well as the cervical premotor circuits the apparent rate of divergence was higher but with a decreased density of labelled premotor neurons. Despite the low proportion of divergent premotor neurons amongst the total premotor population, it is possible that these neurons effectively modulate the synchrony of MN activation and participate in co-activation or co-inhibition of different MN populations.

What proportion of premotor neurons project to more than one motor pool? To investigate the presence of premotor neurons projecting to multiple motor pools in the spinal cord, we used RabV tracing, injecting ΔG-RabV expressing eGFP or mCherry into different pairs of muscles. Although this technique allowed for visualization of divergent premotor neurons throughout the spinal cord, the proportion of divergent premotor neurons has undoubtedly been underestimated. A divergent neuron will be double labelled only if each virus has been efficiently transmitted across its synapses with motoneurons from both motor pools. Therefore, due to the stochastic nature of the process of crossing a synapse, any given transfer efficiency lower than 100% will inevitably give rise to an underestimate of the real number of divergent neurons. The efficiency of trans-synaptic jumps for the SADB19 rabies virus that we used is unknown, and may depend in part on the type of synapse, with stronger connections facilitating transmission of the virus (Ugolini, 2011). The only indirect indication of efficiency comes from the direct comparison of the SADB19 and the more efficient CVS-N2c strains, for which there was at least a 4-fold increase in the ratio of local secondary to primary infected premotor interneurons (Reardon et al., 2016). This result suggests that the trans-synaptic efficiency of SADB19 is no higher than 25%. While there is no evidence for a bias towards stronger or weaker synapses (i.e., the actual number of physical contacts) between proximal and distal premotor interneurons, such a bias could affect efficiency of viral transmission, and could thus also have potentially skewed our relative estimate of divergence. With the simplifying assumption that the efficiencies of viral transfer are equal and independent from each other across spinal cord regions, we simulated a double injection experiment, extracting a binomial distribution, and calculated the relation between the observed and true rate of divergence. With a jump efficiency of 25%, the 4% divergence rate we observed in the lumbar spinal cord would correspond to an actual rate of divergence of 18% (see Figure S6). And this calculated rate is almost certainly an underestimate because of the phenomenon of viral interference, whereby there is a reduced probability of subsequent infection with a second RabV after a window of a few hours after the first infection (Ohara et al., 2009). It is therefore likely that the actual rate of divergence of premotor circuit throughout the cord is substantially higher than we observed. Specifically, it is possible that the vast majority of, if not all, premotor LDPNs innervate more than one motor pool.

### Mapping premotor circuits using the ChAT-Cre;RΦGT mouse

In our experimental model, the rabies glycoprotein is expressed only in neurons expressing ChAT, such as MNs. By restricting primary infection to specific MNs via intramuscular injection of RabV, trans-synaptic viral spread was thus restricted to neurons pre-synaptic to the infected MN population. It is therefore theoretically possible that there might be double jumps via other presynaptic cholinergic neurons such as medial partition neurons (V0_C_ neurons; (Zagoraiou et al., 2009). Motoneurons also form synapses with other MNs (Bhumbra and Beato, 2018), so it could also be possible that specificity is lost due to second order jumps via these cells. We consider double jumps unlikely for two main reasons: 1) following muscle injections, the first transsynaptic labelling occurs after 5-6 days. Since the tissue was fixed 9 days after injections, it is unlikely that many secondary jumps could have occurred in such a brief time window. And 2) most pre-synaptic partners of V0_C_ INs are located in the superficial dorsal laminae (Zampieri et al., 2014), a region in which we did not observe any labelled INs. We are thus confident that the labelled neurons are premotor. We also acknowledge the possibility that some of the labelled premotor cells might originate from tertiary infection originating from secondary infection of synaptically connected MNs (Bhumbra and Beato, 2018). Such events might be rare (Ronzano et al., 2021) and would not alter our findings on the organization of divergent premotor neurons, since we have shown that their distributions are similar, regardless of the particular pair of injected muscles.

### Premotor interneurons innervating antagonist motor pools: implications for movement

The similar rate of divergence between synergist and antagonist pairs might be surprising. But divergence to agonist and antagonist motor pools has been shown in adult mice (Gu et al., 2017), indicating that these circuits are not limited to an early developmental stage. Apart from the cervical divergent premotor LDPNs that are likely to represent a rather homogenous group of excitatory neurons, the divergent premotor neurons in the thoracic and lumbar regions could be comprised of different neural populations, with a mixed population of excitatory and inhibitory neurons. These INs that project to antagonist motor pools could thus be involved in modulating either joint stiffening (excitatory) or relaxation (inhibitory). For example, during postural adjustment and skilled movements, divergent excitatory premotor INs would lead to co-contraction of antagonist muscles to facilitate an increase in joint stiffness and to promote stability (Hansen et al., 2002; Nielsen and Kagamihara, 1993, 1992). In invertebrates, co-contraction of antagonist muscles has also been described preceding jumping (Pearson and Robertson, 1981): co-contraction could thus also be important for the initiation of movement.

On the other hand, divergent inhibitory premotor neurons would lead to joint relaxation. This phenomenon is less well studied (Leis et al., 2000; Manconi et al., 1998). One example could be their involvement in the loss of muscle tone that accompanies rapid eye movement sleep (Uchida et al., 2021; Valencia Garcia et al., 2018). Thus, we provide evidence that the spinal cord contains premotor circuits that could modulate joint stiffness and relaxation.

### Projections of long descending propriospinal neurons to multiple motor pools

In the cat, long descending fibres originating in the cervical cord have been shown to innervate lumbar MNs (Giovanelli Barilari and Kuypers, 1969) and trigger monosynaptic potentials (Jankowska et al., 1974). The existence of LDPNs has been confirmed anatomically in neonatal mice (Ni et al., 2014) and functionally in adult cats (Alstermark et al., 1987a, 1987b), where they are thought to play a role in posture and stability. Our study confirms the existence of premotor LDPNs, and also indicates that they have a high rate of divergence (up to ∼40% compared to ∼13% for thoracic neurons). Most cervical LDPNs are clustered in contralateral lamina VIII, are virtually all excitatory, and have a distinct morphology with somal size ∼2-fold larger than other local cells. These findings contrast with the divergent premotor neurons found in the thoracic spinal cord: these are distributed in ipsilateral lamina VI and VII as well as in contralateral lamina VIII and thus comprise multiple neuronal populations. In contrast to thoracic divergent premotor neurons, cervical LDPNs may be rather homogeneous in function. Given that at least some of these premotor LDPNs project to both lumbar and cervical MNs, it is possible that they also project to motor pools controlling muscles across multiple joints of the same limb. With their apparent widespread divergence, it is possible that these LDPNs are involved in producing widespread increases in muscle tone.

One step towards being able to further assess the function of this population of interneurons would be through understanding their lineage. Although only a subset of labelled LDPNs were clearly V0_V_ derived in our experiments, the poor detection of the Lhx1 transcription factor in postnatal mice (Figure S3), may have limited our sensitivity. Further experiments using a Dbx1-IRES-GFP mouse line (Bouvier et al., 2010), for example, could be done to confirm the identity of these divergent cervical LDPNs. Furthermore, genetic access to this particular set of INs would allow the design of experiments aimed at acute and specific activation or inactivation of divergent LDPNs, and could unravel their anatomy and function in behaviour.

### Concluding remarks

The completion of movements requires well controlled muscle contractions across multiple joints within and between limbs. The control of any one joint is analogous to the production of syllables of speech, with the three most fundamental syllables of movement being a change in joint angle (requiring reciprocal inhibition of flexors and extensor MNs), a stiffening of a joint (requiring co-activation of flexors and extensor MNs), and a relaxation of a joint (requiring co-inhibition of flexor and extensor MNs). While neural circuits for reciprocal inhibition have been well studied over many decades (Eccles, 1969; Eccles et al., 1956), circuits for stiffening or relaxation have not been. Here, we present anatomical evidence using mRabV tracing that demonstrates that such circuits are present within and distributed throughout the spinal cord. Thus, the spinal cord itself has the control mechanisms that lead to the production of the fundamental syllables of movement.

## Materials and Methods

### Mouse strains

All experiments (n=27) were performed according to the Animals (Scientific Procedures) Act UK (1986) and certified by the UCL AWERB committee, under project licence number 70/7621. Homozygous ChAT-IRES-Cre mice (Rossi et al., 2011, Jackson lab, stock #006410) crossed with homozygous RΦGT mice (Takatoh et al., 2013), Jackson lab, stock #024708) were used for double injections (see Table S1 the detail of animal use for each type of injection). For single injections, homozygous ChAT-IRES-Cre mice were crossed with hemizygous GlyT2-eGFP mice (BAC transgene insertion in exon 2 of *Slc6a5* gene allowing specific eGFP expression in GlyT2 positive cells, MGI:3835459, Zeilhofer et al., 2005) and their eGFP positive offspring was mated with homozygous RΦGT (see Table S1).

### Virus production, collection, and titration

We used the glycoprotein G-deleted variant of the SAD-B19 vaccine strain rabies virus (a kind gift from Dr M. Tripodi). Modified rabies virus (ΔG-Rab) with the glycoprotein G sequence replaced by mCherry or eGFP (ΔG-Rab-eGFP/mCherry) was produced at a high concentration with minor modifications to the original protocol (Osakada et al., 2011). BHK cells expressing the rabies glycoprotein G (BHK-G cells) were plated in standard Dulbecco modified medium with 10% fetal bovine serum (FBS) and split after 6-7 hours incubating at 37°C and 5% CO_2_. They were inoculated at a multiplicity of infection of 0.2-0.3 with either ΔG-Rab-eGFP or mCherry virus in 2% FBS, and incubated at 35°C and 3% CO_2_. Plates were then split in 10% FBS at 37°C and 5% CO_2_. After 24h the medium was replaced by 2% FBS medium and incubated at 35°C and 3% CO_2_ for 3 days (virus production). The supernatant was collected and medium was added for another cycle (3 cycles maximum), after which the supernatant was filtered (0.45 µm filter) and centrifuged 2h at 19,400 rpm (SW28 Beckman rotor). The pellets were re-suspended in phosphate buffered saline (PBS) and centrifuged together at 21,000 rpm, 4°C, 4 hours in a 20% sucrose gradient. Pellets of each collection were then re-suspended and stored in 5-10 µl aliquots at −80°C.

Virus titration was performed on BHK cells plated in 10% FBS medium at 1.5 x 10^5^ cells/ml and incubated overnight at 37°C and 10% CO_2_ (growth). The virus was prepared for 2 serial dilutions with 2 different aliquots and added in the well after an equal volume of medium had been removed (serial dilution from 10^-3^ to 10^-10^) and incubated 48h at 35°C and 3% CO_2_. The titre was determined from the count of cells in the higher dilution well and was between 10^9^ and 10^10^ infectious units (IU)/ml.

### Intramuscular injections

A subcutaneous injection of analgesic (carprofen, 1 µl, 10% w/v) was given to the neonatal pups (P1-P3) prior to surgery and all procedures were carried out under general isoflurane anaesthesia. After a skin incision to expose the targeted muscle, the virus (1µl) was injected intramuscularly using a Hamilton injector (model 7652-01) mounted with a bevelled glass pipette (inner diameter 50-70 µm). The mice were injected in tibialis anterior (TA) and peroneus longus (PL) (ankle flexor pair), lateral and medial gastrocnemius (LG and MG) (ankle extensor pair) for synergist pairs and TA and LG for antagonist pairs. In hindlimb/forelimb double injections, the LG and MG were both injected with 1 µl of one RabV to increase the number of long projecting cells infected. In addition, 1 µl of the second RabV was injected in forearm muscles (FMs, see Table S2) without selecting a specific muscle. The injected viruses were used at a titre between 10^9^ and 10^10^ IU/ml. The incisions were closed with vicryl suture, and the mice were closely monitored for 24 hours post-surgery. Mice were perfused 9 days after the injections. Following perfusion, the specificity of injection was checked on the injected limb when synergist pair of muscles had been injected.

### Tissue collection and immunohistochemistry

The mice were perfused with phosphate buffer solution (PBS, 0.1 M) followed by PBS 4% paraformaldehyde under terminal ketamine/xylazine anaesthesia (i.p. 80 mg/kg and 10 mg/kg respectively). The spinal cords were then collected through a ventral laminectomy and post-fixed for 2 hours. The cords were divided into the different parts of the spinal cord (cervical (C1-T1), thoracic (T2-T11), lumbar (L1-L6) and sacral (S1-S4)), cryoprotected overnight in 30% sucrose PBS, embedded in optimal cutting temperature compound (Tissue-Tek) and sliced transversally (30 µm thickness) with a cryostat (Bright instruments, UK). Sections were incubated with primary antibodies for 36h at 4°C and with secondary antibodies overnight at 4°C in PBS double salt, 0.2% Triton 100-X (Sigma), 7% donkey normal serum (Sigma). The primary antibodies used were: goat anti-choline acetyltransferase (ChAT, 1:100, Millipore, AB144P), chicken anti-mCherry (1:2500, Abcam, Ab205402), rabbit anti-GFP (1:2500, Abcam, Ab290), guinea pig anti-vGluT2 (1:2500, Millipore, AB2251-I), and rabbit anti-Lhx1 (1:5000, from Dr. T Jessell, Columbia University, New York); and the secondary antibodies: donkey anti-rabbit Alexa 647 (1:1000, Abcam, Ab150079), donkey anti-goat preadsorbed Alexa 405 (1:200, Abcam, Ab175665), donkey anti-rabbit Alexa 488 (1:1000, Thermofisher, A21206), and donkey anti-chicken Cy3 (1:1000, Jackson ImmunoResearch, #703-165-155). The slides were mounted in Mowiol (Sigma, 81381-250G) and coverslipped (VWR, #631-0147) for imaging.

### Confocal imaging and analysis

Images of the entire sections were obtained using a Zeiss LSM800 confocal microscope with a 20x air objective (0.8 NA) and tile advanced set up function (ZEN Blue 2.3 software). A 63x oil objective was used for Airy scan imaging of somata and excitatory boutons. Tiles were stitched using Zen Blue and analyses were performed using Zen Blue and Imaris (Bitplane, version 9.1) software packages. Location maps were plotted setting the central canal as (0,0) in the (x,y) Cartesian system and using the “Spots” function of Imaris. The y-axis was set to the dorso-ventral axis. Positive values were assigned for dorsal neurons in the y-axis and ipsilateral (to the hindlimb injection) neurons in the x-axis. Coordinates were collected on every section and normalized through the cervical, thoracic, lumbar and sacral parts separately using grey matter borders and fixing the width and the height of the transverse hemisections. To calculate divergence rates, in the lumbar cord all infected premotor INs (eGFP+, mCherry+ and eGFP+mCherry+) were quantified in 1 of every 3 sections to avoid counting the same cells twice on consecutive sections. In the cervical, thoracic and sacral regions, all cells were quantified, as their low density allowed for manually excluding premotor INs found in consecutive sections.

### Statistics

All statistical analyses and plots were made using R (R foundation for statistical computing, Vienna, Austria, 2005, http://www.r-project.org, version 3.6.2) and GraphPad PRISM (version 7.0). To compare cell sectional areas, non-parametric rank tests were used as specified in each related result. The numbers of animals/cells in each experiment and statistical tests used are reported in the figure legends or directly in the text. Results and graphs illustrate the mean ± standard deviation. Statistical significance levels are represented as follows: * p < 0.05, ** p < 0.01, *** p < 0.001, **** p < 0.0001 and ns: not significant.

## Acknowledgments

We are grateful to Dr N. Zampieri and all the members of Brownstone and Beato labs for insightful comments on the manuscript. We want to thank S. Morton for providing Lhx1 antibodies and Prof L. Greensmith for access to her lab facilities. This work was supported by a Leverhulme Trust grant (grant number RPG-2013-176) and a BBSRC grant (BB/L001454) to MB and a Wellcome Trust Investigator Award to RMB (110193). RMB is supported by Brain Research UK.

## Declaration of interests

The authors declare no conflict of interest.

---

**Figure S1:**
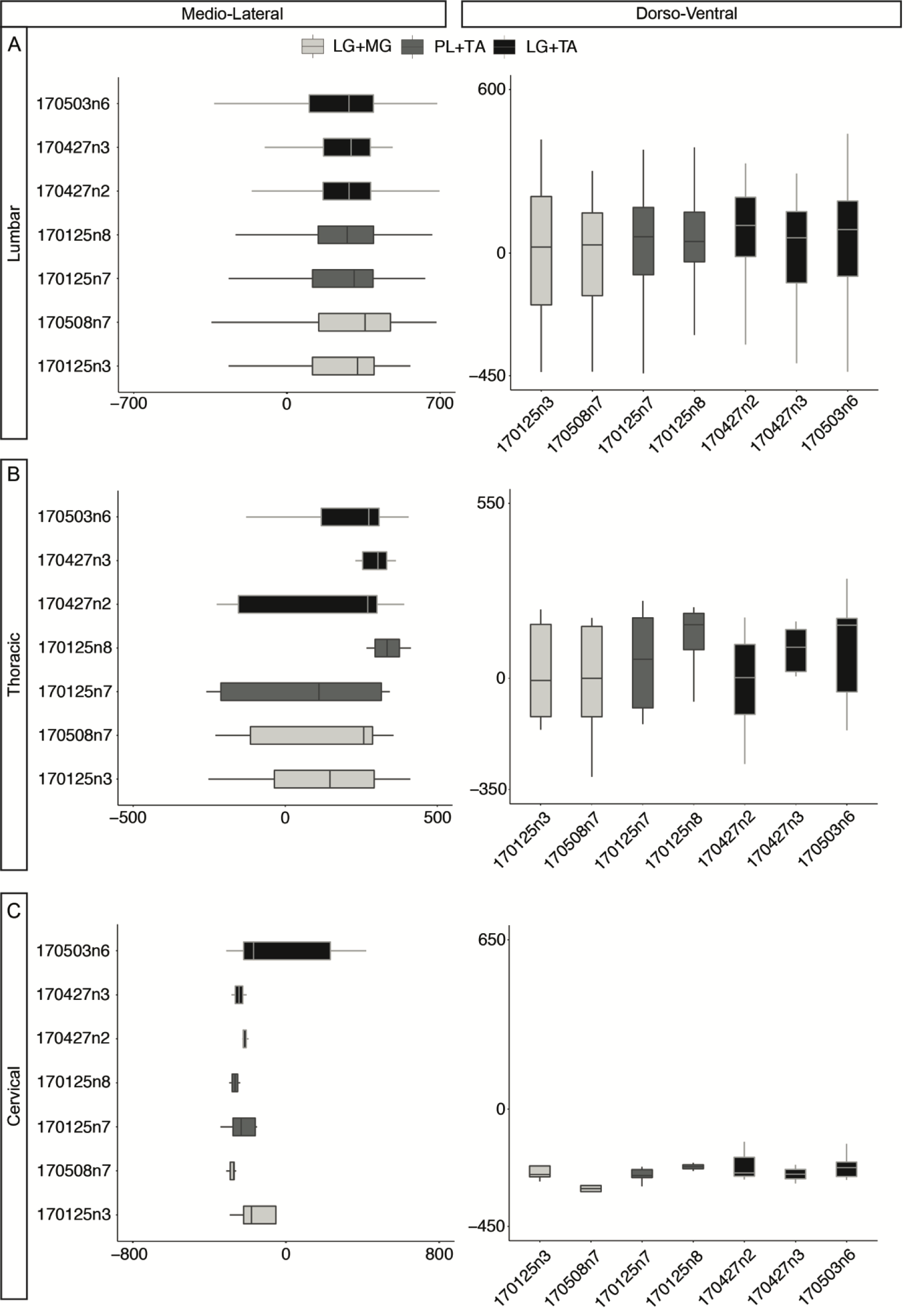
Medio-lateral and dorso-ventral distributions of divergent premotor neurons across individual experiments. Boxplot showing the medio-lateral and the dorso-ventral distributions of divergent premotor neurons in the (A) lumbar, (B) thoracic and (C) cervical parts across individual experiment. The distributions are consistent across experiment and pairs of muscles injected.

**Figure S2:**
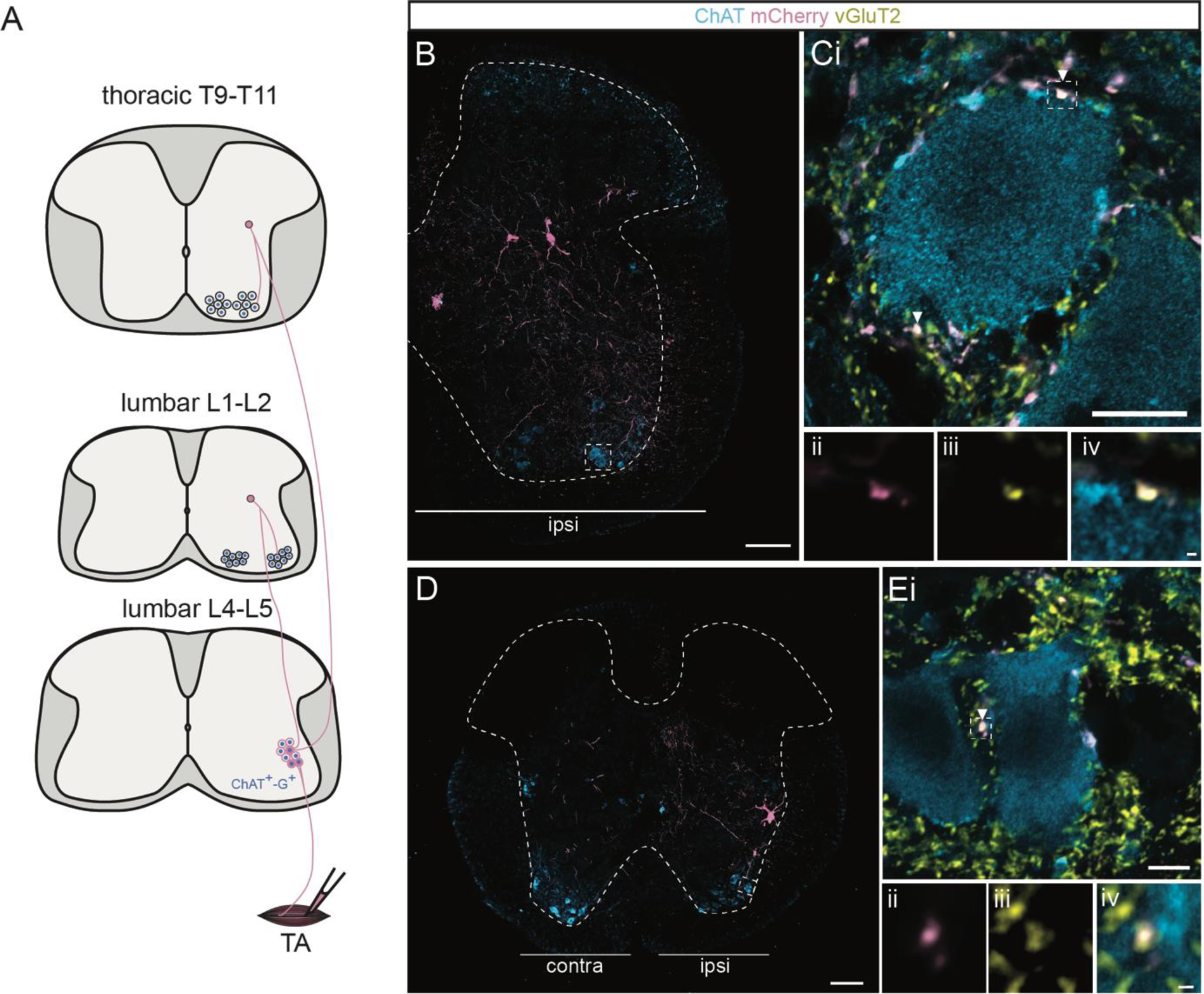
Excitatory boutons on MNs belonging to infected premotor neurons, reveal divergence through different segments and parts of the cord. **(A)** Schematic showing experimental design and projections visualized thanks to the study of excitatory boutons apposed on MNs. **(B, D)** Representative image of a traverse section in the (B) L1 segment (upper lumbar) and (D) thoracic part, following an injection of ΔG-Rab-mCherry in the TA, showing ChAT (blue grey) and mCherry (pink). MNs with vGluT2+ (yellow); mCherry+ boutons affixed on them are highlighted in the dashed boxes. The dashed contours drawn follows the grey matter contours. **(C, E)** Dashed box areas at higher magnification. Arrowheads show vGluT2+; mCherry+ boutons on (C) L1 and (E) thoracic MNs. Dashed boxes highlight the boutons showed at higher magnification in (Cii-iv and Eii-iv). Scale bars: (B, D) 100 µm; (Ci,Ei) 10 µm; (Civ, Eiv) 0.5 µm.

**Figure S3:**
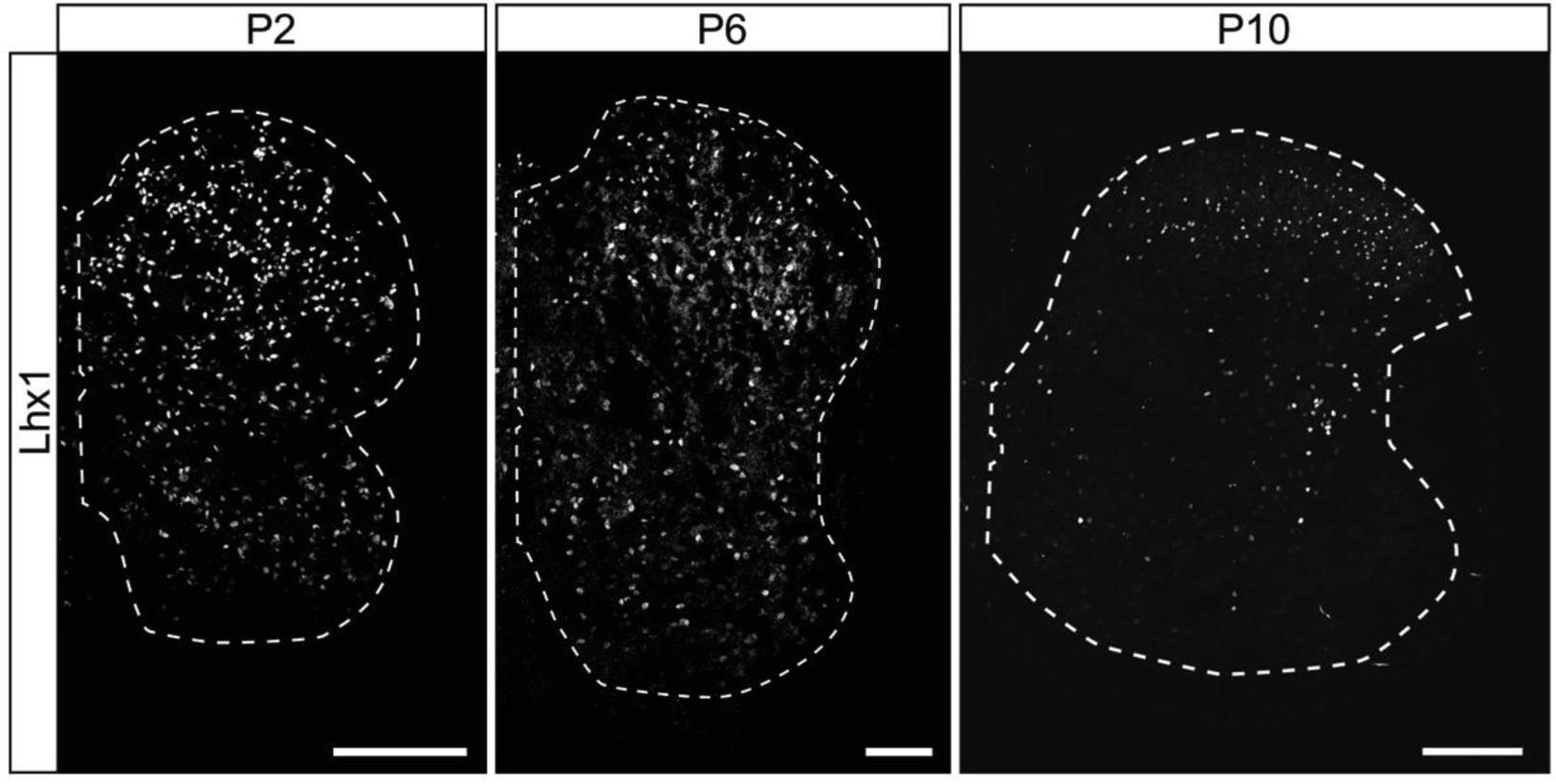
The number of neurons labelled with anti-Lhx1 antibody decreases in the spinal cord over postnatal development. Lhx1 staining at P2, P6 and P10 showing the decrease in the number of Lhx1 positive cells along postnatal development of the spinal cord. The dashed contours drawn follows the grey matter contours. Scale bars: 100 µm.

**Figure S4:**
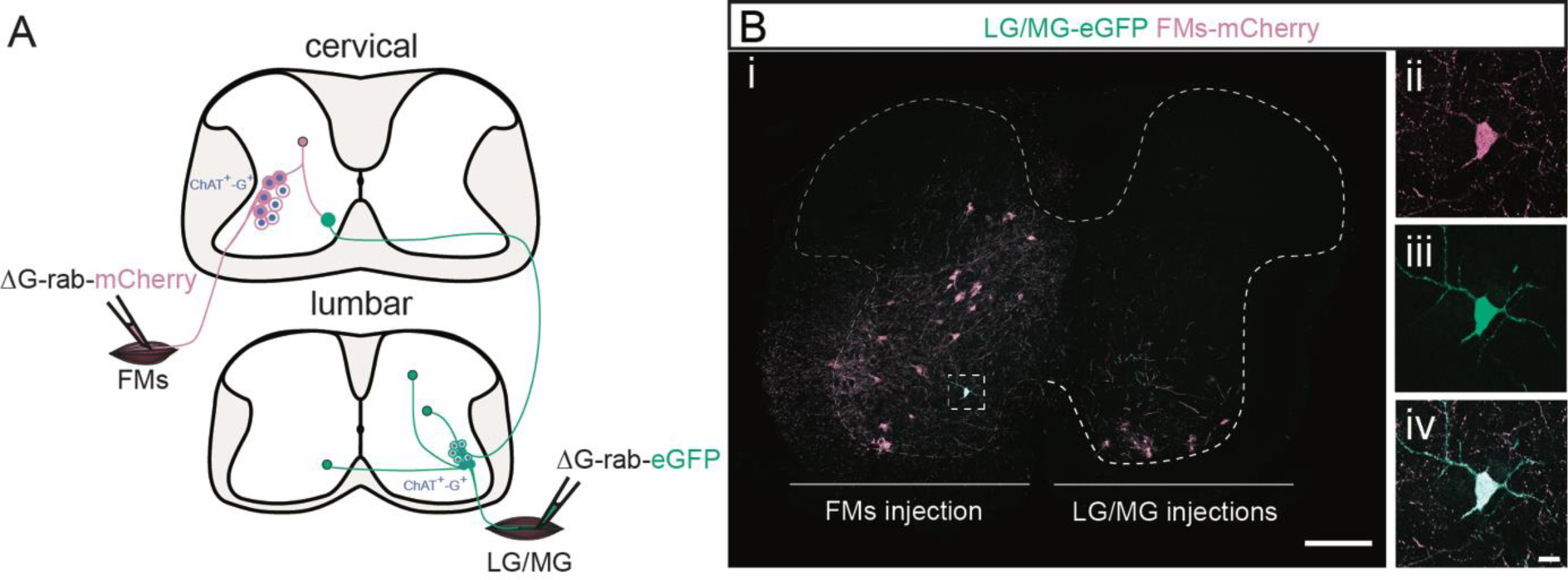
Cervical premotor LDPNs innervating ipsilateral cervical and contralateral lumbar MNs exist. **(A)** Experimental strategy to determine whether divergent cervical premotor LDPNs that innervate ipsilateral cervical and contralateral lumbar motor pools do exist. **(Bi**) Representative example of a transverse section from the cervical cord following an injection in contralateral GS (ΔG-Rab-eGFP) and FMs (ΔG-Rab-mCherry) muscles, showing a premotor LDPN infected from the two injections. The dashed box highlights the divergent premotor LDPN. The dashed contour drawn follows the grey matter contour. **(ii-iv)** Dashed box area at higher magnification. Scale bars: (Bi) 200 µm; (Biv) 20 µm.

**Figure S5:**
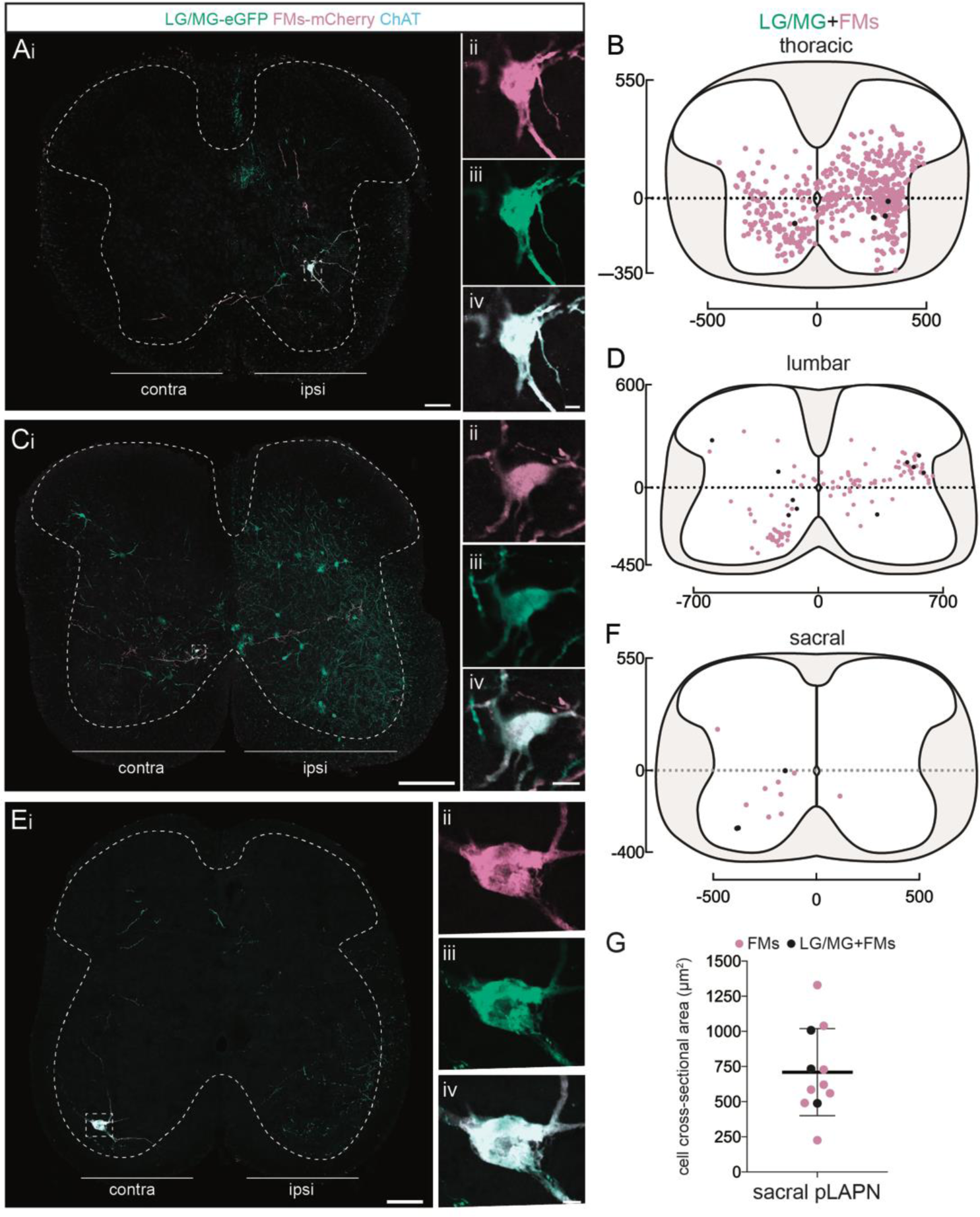
Premotor LAPNs distribution throughout the spinal cord and divergence to ipsilateral lumbar and cervical MNs. **(Ai, Ci, Ei)** Representative examples of transverse sections from the (A) thoracic, (C) lumbar and (E) sacral cord following injections in homolateral GS (ΔG-Rab-eGFP) and FMs (ΔG-Rab-mCherry) showing ChAT (blue grey), GFP (green) and mCherry (pink). Dashed boxes highlight INs that were infected from the injections in homolateral GS and FMs. The dashed contours drawn follow the grey matter contours. **(Aii-iv, Cii-iv, Eii-iv)** High-magnification of the previous boxes showing double infected premotor neurons in the (Aii-iv) thoracic, (Cii-iv) lumbar and (Eii-iv) sacral cord. **(B, D, F)** Distributions of the infected ascending (pink) and divergent (black) premotor neurons in the (B) thoracic, (D) lumbar and (F) sacral cord following injections in homolateral GS (ΔG-Rab-eGFP) and FMs (ΔG-Rab-mCherry). **(G)** Plot showing the sectional area of the sacral premotor LAPNs infected from GS and FMs injections (black) or from the injection in FMs only (pink). Scale bars: (Ai, Ei) 100 µm; (Ci) 200 µm (Aiv, Civ, Eiv) 10 µm.

**Figure S6:**
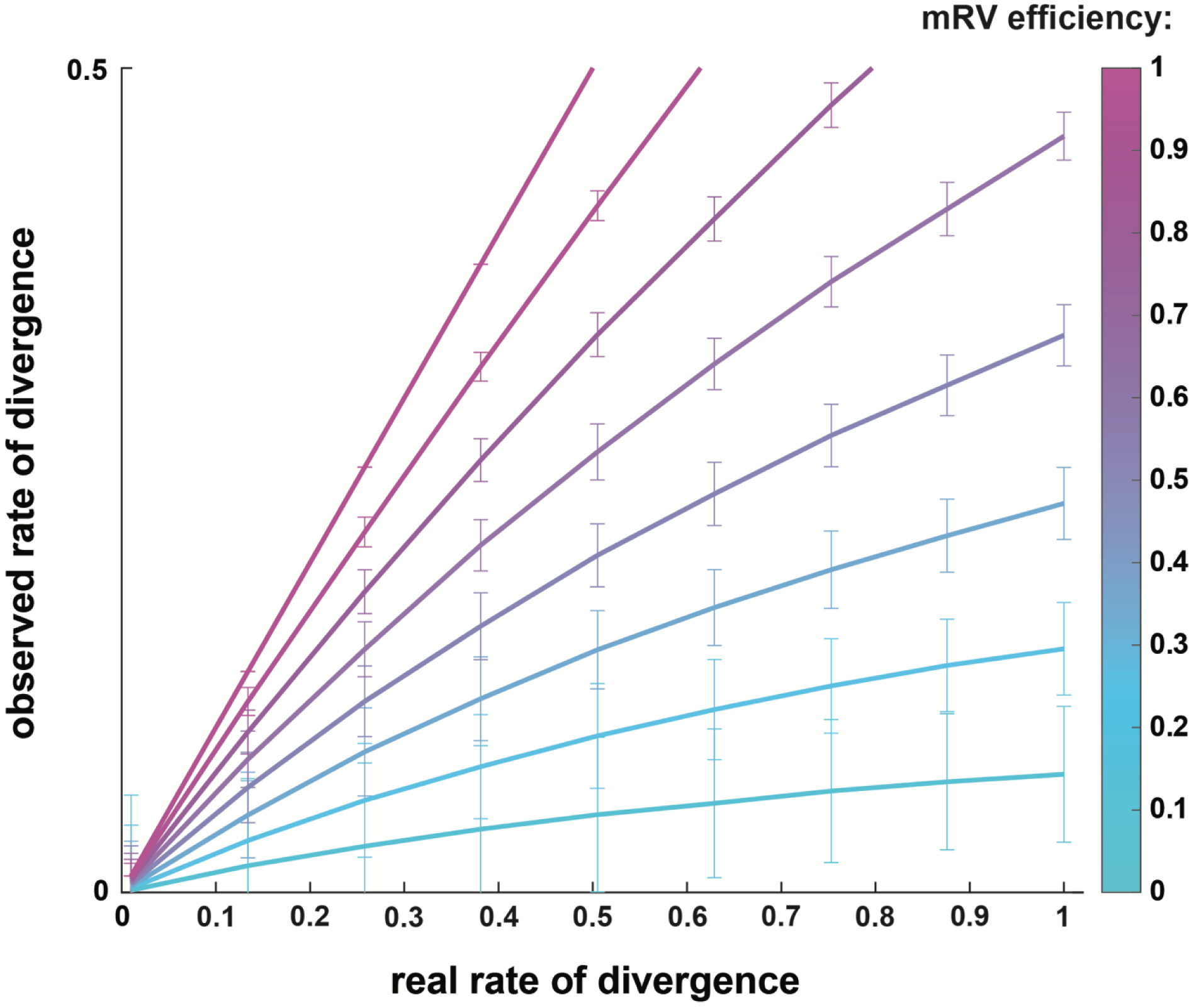
Simulation comparing observed vs real rates of divergence depending on transsynaptic mRV efficiency. Simulation of the spreading of mRV in premotor circuits following double injections, extracted from a binomial distribution. Plot showing the relation between observed rate of divergence depending on the real rate of divergence within premotor spinal circuits. This simulation was run with the simplifying assumption that the efficiencies of viral transfer are equal and independent from each other across spinal cord regions.

**Table S1:** Details of the mouse used for each experiment.

| Figure in which samples are used | Mouse strain | Muscles injected | Number of animals | Used for which study |
| --- | --- | --- | --- | --- |
| 1, 2, 3, S1 | ChAT-Cre;RΦGT | LG, MG | 2 | this study; Ronzano et al, 2021 |
| 1, 2, 3, S1 | ChAT-Cre;RΦGT | TA, PL | 2 | this study; Ronzano et al, 2021 |
| 1, 2, 3, S1 | ChAT-Cre;RΦGT | TA, LG | 3 | this study; Ronzano et al, 2021 |
| 4 | ChAT-Cre;RΦGT ;GlyT2-eGFP | LG | 3 | this study only |
| 5 | ChAT-Cre;RΦGT | LG+MG | 4 | this study only |
| 6, S5 | ChAT-Cre;RΦGT | LG+MG, FMs homolateral | 6 | this study only |
| S2 | ChAT-Cre;RΦGT | TA | 1 | this study only |
| S3 | ChAT-Cre;RΦGT | NA | 3 | this study only |
| S4 | ChAT-Cre;RΦGT | LG+MG, FMs contralateral | 3 | this study only |

**Table S2:** Details of muscles infected following forearm injections. (+) means that a fluorescent signal was found in muscle fibers, (-) means that no fluorescence was observed following muscle dissections.

| Sample | Type of injection | Extensor carpi radialis longus | Extensor carpi radialis brevis | Flexor carpi radialis | Extensor carpi ulnaris | Flexor carpi ulnaris | Palmaris longus | Digitorum profundus |
| --- | --- | --- | --- | --- | --- | --- | --- | --- |
| S_4_0106 | homolateral | - | - | - | - | - | - | - |
| S_5_0106 | homolateral | - | + | - | - | + | - | - |
| S_6_0106 | homolateral | - | + | - | + | + | - | - |
| S_5_2306 | homolateral | + | + | - | + | + | - | - |
| S_7_2306 | homolateral | + | + | + | - | + | + | + |
| S_8_2306 | homolateral | + | + | - | - | + | - | - |
| S_7_0106 | contralateral | - | - | - | - | + | - | - |
| S_8_0106 | contralateral | + | + | - | + | + | - | - |
| S_10_0106 | contralateral | + | + | - | - | + | - | - |

**Table S3:** Medio-lateral and dorso-ventral distributions of divergent premotor neurons across individual experiments. Distribution of divergent premotor neurons per part of the spinal cord across individual experiments, expressed as median ± first/third quartile.

| Sample | Muscles injected | Lumbar |  | Thoracic |  | Cervical |  |
| --- | --- | --- | --- | --- | --- | --- | --- |
|  |  | Medio-Lateral | Dorso-Ventral | Medio-Lateral | Dorso-Ventral | Medio-Lateral | Dorso-Ventral |
| 170125n3 | LG - MG | 323 ± 95/400 | 22 ± -191/209 | 147 ± -66/300 | -7 ± -128/177 | -180 ± -269/168 | -251 ± -271/-159 |
| 170508n7 | LG - MG | 357 ± 127/474 | 30 ± -157/148 | 258 ± -124/289 | 0 ± -121/167 | -273 ± -301/-265 | -305 ± -317/-264 |
| 170125n7 | PL - TA | 307 ± 112/396 | 60 ± -82/169 | 111 ± -222/316 | 60 ± -96/197 | -234 ± -298/-152 | -255 ± -264/-225 |
| 170125n8 | PL - TA | 275 ± 140/396 | 42 ± -34/150 | 335 ± 294/376 | 169 ± 73/205 | -266 ± -296/-239 | -221 ± -237/-206 |
| 170427n2 | LG - TA | 284 ± 164/385 | 101 ± -14/206 | 272 ± -186/333 | 2 ± -126/108 | -221 ± -226/-195 | -245 ± -270/-125 |
| 170427n3 | LG - TA | 293 ± 160/386 | 56 ± -128/152 | 305 ± 243/349 | 98 ± 14/166 | -245 ± -284/-206 | -249 ± -285/-214 |
| 170503n6 | LG - TA | 284 ± 102/397 | 86 ± -89/191 | 275 ± 86/312 | 167 ± -45/193 | -169 ± -228/247 | -225 ± -266/-187 |

## Notes

### Competing Interest Statement

The authors have declared no competing interest.

### Summary of Updates

Correction of minor mistakes in the number of reported infected cells

